# High Throughput Screening at the Membrane Interface Reveals New Inhibitors of Amyloid-β

**DOI:** 10.1101/853499

**Authors:** Sarah J. Cox, Brian Lam, Ajay Prasad, Hannah A. Marietta, Nicholas V. Stander, Joseph G. Joel, Bikash R. Sahoo, Fucheng Guo, Andrea K. Stoddard, Magdalena I. Ivanova, Ayyalusamy Ramamoorthy

## Abstract

Amyloid-β aggregation at the cell-membrane of neruonal cells is implicated as a source of toxicity for Alzheimer’s disease. Small molecules have been studied for their ability to supress amyloid aggregation and toxicity, but the presence of membranes negate their activity. Here, we have identified 5 small molecules that are active at the membrane interface.

---

Alzheimer’s disease (AD) is a deadly and debilitating ailment that currently affects 50 million people worldwide.^1^ Early research into AD focused on the presence of indicative protein amyloid-beta (Aβ) fibrils due to their prominence in postmortem examination of patients’ brains. However, it is now hypothesized that small, toxic, intermediate species, known as oligomers, are the predominant toxic amyloid-beta (Aβ_40_) species in AD.^2^ Aβ peptides are produced from the cleavage of amyloid precursor protein (APP) in the extracellular membrane by β and γ-secretases. Cellular membranes have been implicated to be a site of potential toxicity and can act as a catalyst for amyloid aggregation.^3^ Some oligomers are proposed to impart their toxic function by interacting directly with the cell membrane of neurons then disrupting and permeabilizing the membrane. As a result, non-selective ion channels and large pores are created which, in turn, ablate the charge gradient necessary for neuronal function.^4^ Many studies suggest that lipid membranes are able to accelerate the aggregation of Aβ as well as facilitate the formation of unique structures of Aβ species that are specific to lipid bilayer disruption.^5^

There has been extensive investigation into small molecules with the ability to modulate the aggregation of Aβ in solution.^6^ However, the search for modulators of Aβ aggregation has relied heavily on serendipity; often times, a novel class of inhibitors is accidentally discovered, and improved analogues are subsequently synthesized.^7^ Relying on accidental discoveries is unlikely to generate a diverse enough chemical portfolio to successfully generate a drug candidate that can demonstrate clinical efficacy. Thus, it is essential to identify new and novel chemical species which may be specifically capable of modulating membrane-assisted Aβ_40_ aggregation for use as toxic Aβ oligomer probes.^2^ Here, through the usage of a small molecule library, 5 compounds have been identified that modulate the formation of Aβ_40_ aggregates in the presence of lipid membrane. These small molecules represent an avenue for the development and further investigation of Aβ_40_ and membrane interactions.

Using a library of over 1,800 compounds, selected for their chemical diversity and biological activity, the screening was performed by the addition of biologically obtained Aβ_40_ in the presence of large unilamellar vesicles (LUVs) composed of a mixture of 7:3 molar ratio of DOPC:DOPG, which represents the charge distribution of eukaryotic membranes (Figure 1A). To probe the interactions between the lipid bilayer, small molecules, and Aβ_40_, we used a fluorescence readout assay regularly employed in amyloid studies using Thioflavin-T (ThT) dye. ThT assays provide insights into the kinetics of amyloid formation, which facilitates the identification of compounds that are able to inhibit the formation of β-sheet rich amyloid aggregates. Signal intensity can be proportional to the fibers present and decreases in intensity can be indicative of a decrease in overall fiber content (Figure S1). We optimized screening conditions, achieving a Z-score of 0.46 (Figure 1C). 40 reproducible hits were selected after ruling out initially fluorescent compounds and compounds which did not give a matching read-out in twin sets of plates. These 40 compounds were then used for a concentration response curve (CRC) titration screen to determine the activity of the compounds in a range of concentrations (Figures S2 and S3). Results from the CRC screen helped us to narrow down the 40 initial hits to 21 primary hits based on the calculated IC50 values and the exclusion of known compounds with PAINS properties (Figure 1D, Tables S1 and S2).^7^

**Figure 1.**
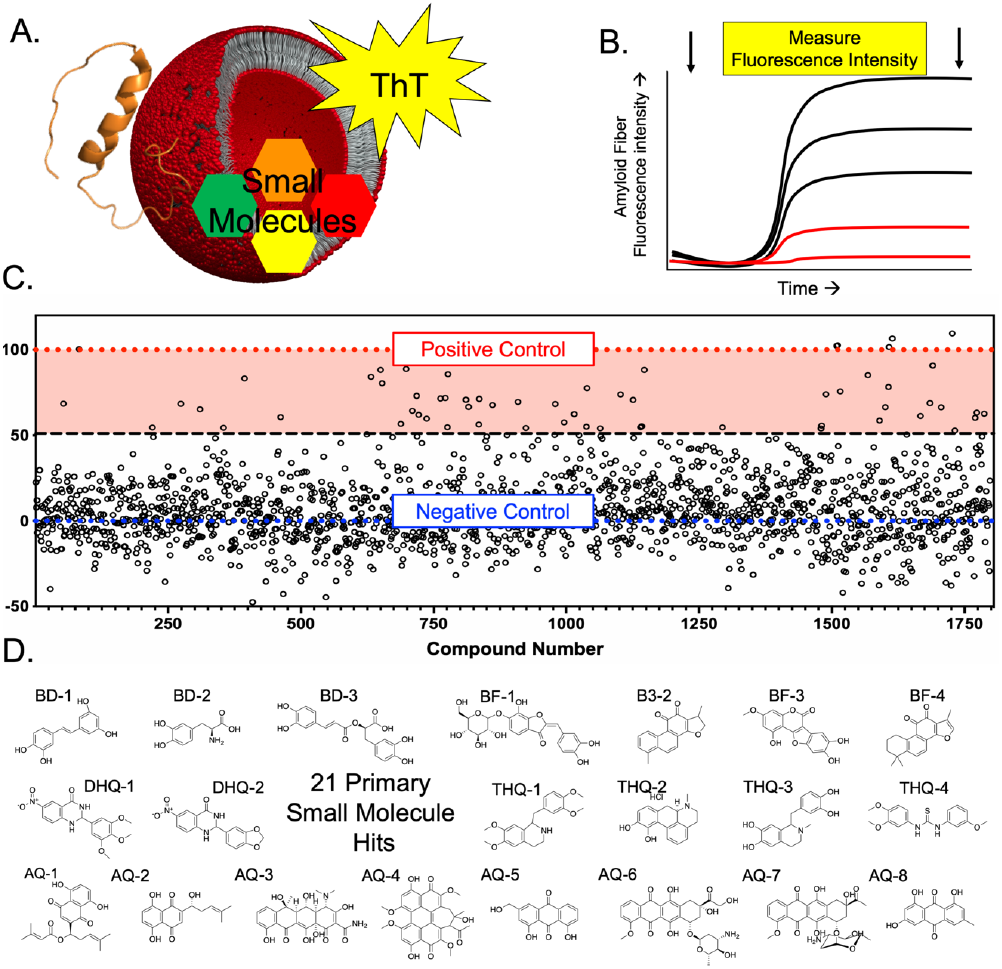
(A) An illustration of the components of the screen: Aβ_40_ monomer (2LFM), LUVs, small molecules and ThT. (B) schematic of reading the assay plates before aggregation and then heating and shaking the plates before reading the final fluorescence intensity after 24 hours of incubation. (C) Final fluorescent intensity of every compounded screened for inhibition. Value of twin plates are averaged and normalized in respect to the positive and negative controls. (D) 21 primary small molecule hits chosen after initial screen and CRC testing.

The selected 21 compounds were initially subjected to full ThT kinetic profiles with measurements taken every 5 minutes (Figure S4). Each compound was tested at 10, 5 and 2 molar equivalencies in respect to the concentration of Aβ_40_ while in the presence of 500 μM LUVs. Initially by the ThT assay, many appear to be promising and robust inhibitors, with many compounds negating the aggregation fully at all the concentrations tested. However, while as critical as the ThT assay is to studying amyloid aggregation, it is also subject to fluorescent quenching, overlap, or displacement by other compounds. Because of this, secondary confirmation not relying on fluorescence was performed to further narrow down the hit compounds. To do this we used the dot blot assay utilizing the OC anti-amyloid fiber antibody, which is known to bind to the general amyloid fiber β-sheet epitope (Figures 2A and S5).^9^ While many of the compounds looked to be complete inhibitors by the ThT assay, the strong antibody reactivity observed for Aβ in presence of these compounds indicated that they do not inhibit fiber formation. After identifying compounds that interfere with ThT, we then examined the compounds that gave 50% or less reactivity by the dot blot assay by examining them via Transmission Electron Microscopy (TEM) (Figure 2B). Out of the 15 compounds investigated, 5 of them inhibited Aβ_40_ fibers, thus AQ-4, THQ-1, BF-3, DHQ-1 and DHQ-2, which were selected for a deeper investigation. Of the other 10 compounds that had fiber formation identified at this stage, many of them exhibited very interesting and distinct fiber morphologies, which could be of interest for further investigation, as some of them have been reported to be amyloid inhibitors in the absence of membrane.

**Figure 2.**
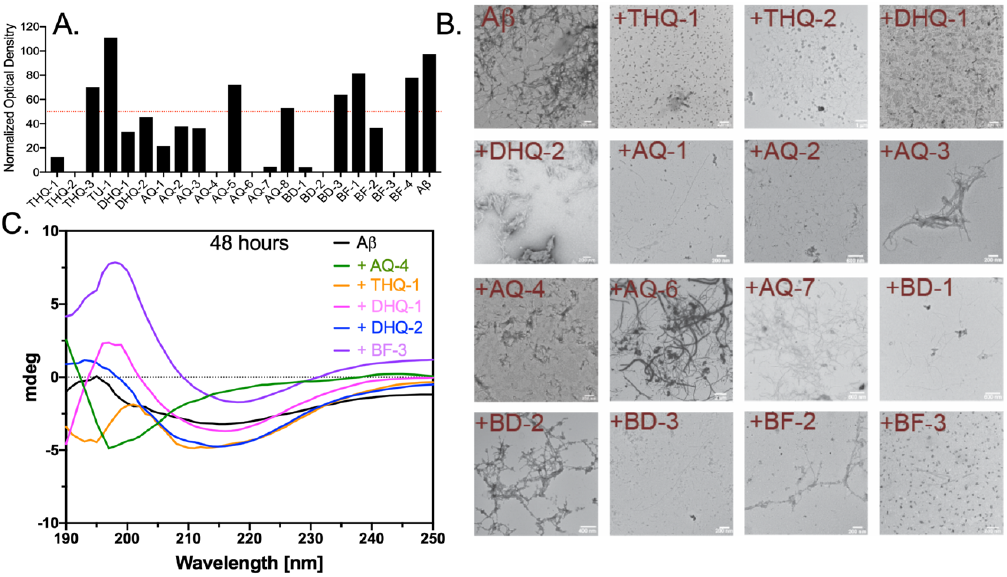
(A) Signal intensities from dot blot assay using the OC antibody. Samples were measured after ThT experiments with 5 equivalents of a compound with respect to Aβ_40_. (B) TEM images for compounds that gave less than 50% antibody reactivity. (C) CD spectra of 25 μM Aβ_40_ in the presence of 500 μM of LUVs with 50 μM loaded compound after 48 hours of incubation with background subtraction of loaded LUVs.

Understanding the secondary structure transitions is important for amyloid investigation, as there is a known shift from a random-coil monomer to β-sheet fiber. To study the compounds’ effects on Aβ secondary structure, the 5 nonwater-soluble compounds were incorporated in the lipid bilayer, which was confirmed by UV-Vis (Figure S6). Upon incorporation of the compounds in the lipid bilayer at a 10:1 lipid to compound molar ratio, we monitored the Aβ structural transitions by circular dichroism (CD) experiments as well as by ThT kinetics (Figures 2C, S7, S8). After 24 hours, AQ-4 and BF-3 still exhibited a random-coil structure. DHQ-2 and THQ-1 showed a minor helical conformation, while DHQ-1 showed a strong β-sheet conformation for Aβ. Up to 7 days, AQ-4 maintained random-coil conformation, whereas BF-3, DHQ-1, and DHQ-2 showed β-sheet conformation. THQ-1, however, showed poor signal and showed some slight β-sheet characteristics for Aβ but was not fully interpretable at 48 hours, but appeared more clearly β-sheet after 7 days.

Using NMR, we investigated the interaction of ^15^N-labeled-Aβ_40_ with the reported compounds both with and without the presence of loaded LUVs using 2D ^1^H/^15^N SOFAST-HMQC experiments (Figures 3, S9, and S10). This experiment is useful in determining the level of peptide aggregation (or monomer depletion) in solution. Because large aggregates such as amyloid fibers tumble slowly on the NMR timescale they do not contribute to the observed signal and any signal is conferred to be fast tumbling oligomers or monomers. The volume of each peak as well as the signal-to-noise ratio were analyzed (Figures 3 and S11-S13). In the presence of loaded vesicles, Aβ_40_ showed 15 well resolved peaks at time zero, with only 4 peaks with poor S/N were observable after 96 hours. For three of the compounds (DHQ-1, DHQ-2 and BF-3), the observed NMR resonances was found to be distributed throughout the amino acid sequence of Aβ_40_ at time zero and after 24 hours, which was also the case in the samples without lipids. For AQ-4, 5 peaks from the C terminus of Aβ_40_ were seen even after 96 hours of incubation, which was also seen in the sample without lipids, indicating some sort of conserved similarities which could signify that the compound is both interacting with the lipid bilayer as well as Aβ_40_ itself. A possible explanation to this is that the N terminus is bound inside an oligomer or to the membrane with a solvent exposed C terminus tail. Spectra of Aβ with THQ-1 and lipids showed very little signal intensity at time zero, with no visible peaks after 96 hours. The opposite was seen in the THQ-1 sample without lipids, in which well resolved peaks of Aβ_40_ were seen at both time zero and at 96 hours. This indicates that ThQ1 interacts and inhibits Aβ_40_ aggregation but does not interact well in presence of lipids, at least when loaded with lipids, since it appeared to be a strong inhibitor in the assays prior to loading the compounds in LUVs. It is also probable that some of the signal loss and poor S/N could be due to Aβ_40_ binding to 100 nm LUVs that decreases the tumbling rate of LUVs. Experiments utilizing smaller membrane mimetic such as nanodiscs and implementing paramagnetic quenching NMR experiments could be an enlightening next step to understand these systems.

**Figure 3.**
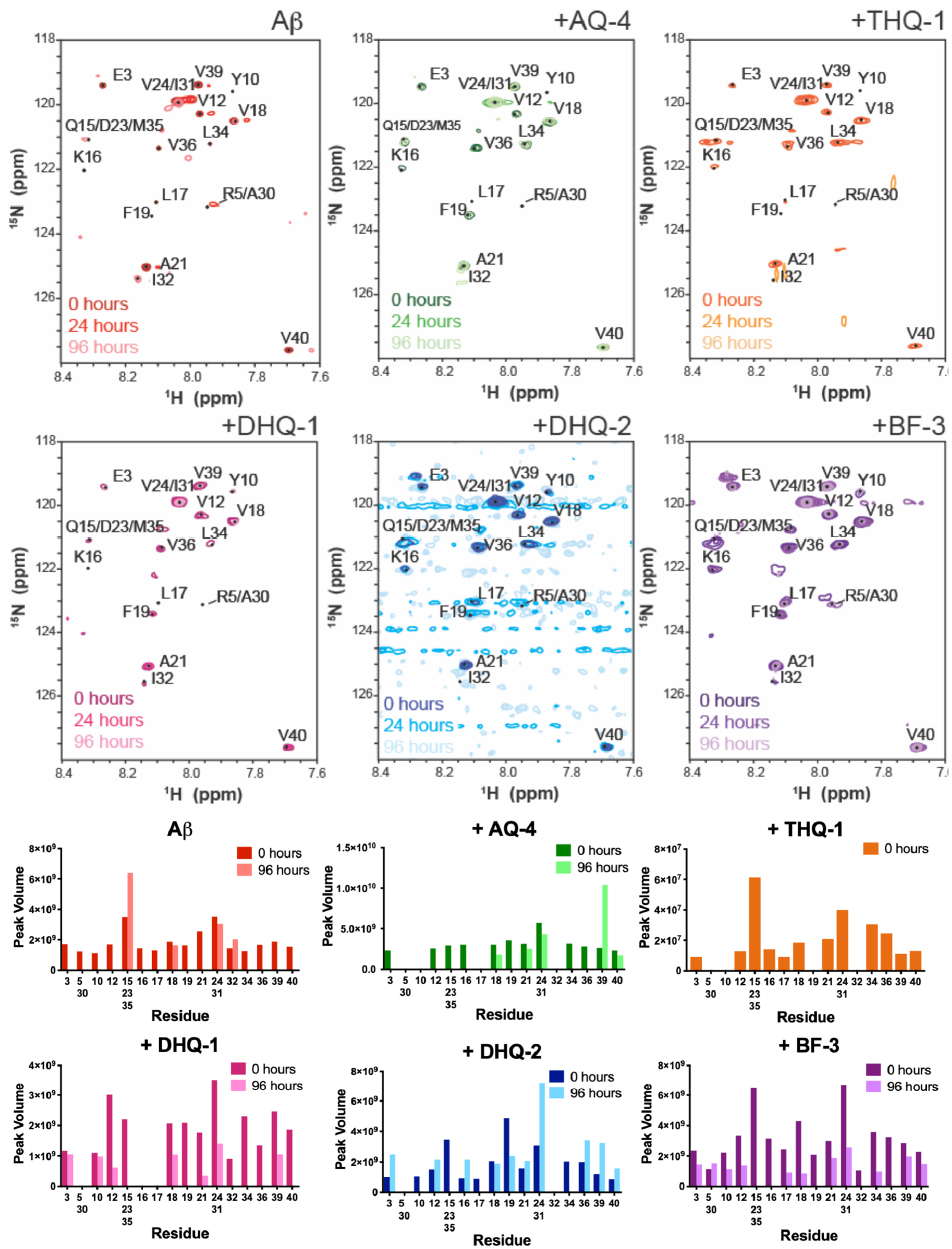
(top 2 rows) SOFAST-HMQC NMR spectra of 25 μM ^15^N-labeled-Aβ_40_ in the presence of 500 μM of LUVs with a 50 μM loaded compound at 0, 24 and 96 hours. (bottom 2 rows) Peak volume of visible peaks at 0 and 96 hours.

After the NMR measurements, the samples in presence of lipids were analyzed using Size Exclusion Chromatography (SEC) (Figures 4 and S14). Three distinct peak areas were seen and the area under each curve was quantified to determine the distribution of aggregates within the sample. Peak area 1 (between 5 and 8 mL) consisted of multiple peaks most likely made up of LUVs and Aβ_40_ amyloid fibers. Peak area 2 (15 mL) most possibly corresponds to an oligomer of 4 monomers (~17 kDa) or an oligomer of 3 monomers with each bound to 2 compounds (~16 kDa). Lastly, Peak area 3 could correspond to monomer and dimer of Aβ_40_, eluting at 21.5 and 20 mL respectively. Since, these samples contain lipids, it is possible that a portion of the intensity of the peaks is also lipids that have been fragmented from the LUVs. For AQ-4, DHQ-1 and BF-3, over 50% of the eluted sample was either in peak area 2 or 3, indicating the presence of a small amount of fibers. While the control, THQ-1, and DHQ-2 samples had 60% or more of the total signal in peak area 1. The control samples showed no signal inside of peak area 2. Additionally, there was a very small peak seen at 11 mL which could indicate a very large oligomer or protofibers. Inside peak area 1, three distinct peaks were observed: one for LUVs at 7.5 mL, one for fibers at 6.5 mL, and in between them; there is also a less resolved peak that may be Aβ_40_ bound to LUVs, which is also not well resolved with the control sample of Aβ_40_ in LUVs. For the control LUVs sample, a small percentage of the total sample eluted in peak area 3, which could correspond to small lipid micelles. This could indicate that some of the signal in the other samples may also contain micelles or lipids that have been fragmented from the bilayer as the result of the peptide aggregation on the membrane.

**Figure 4.**
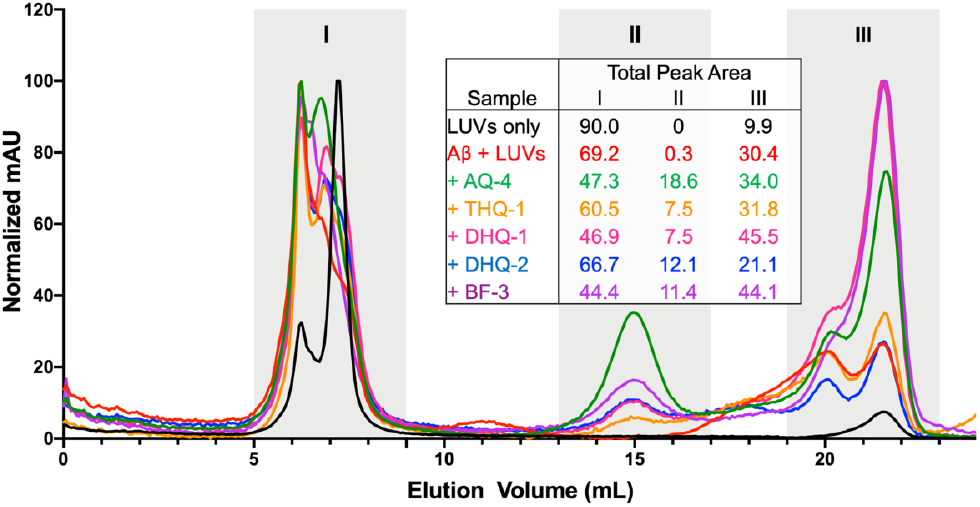
Size exclusion chromatography of the indicated NMR samples and normalized area under the curve of each peak area measured for all samples.

## Conclusions

A high-throughput screen has led to the identification of 5 membrane active Aβ40 amyloid inhibitors, with a brief summary in Table 1. Among them, DHQ-1, DHQ-2 and THQ-1 were found to be the least robust as shown by NMR, SEC and CD results, whereas BF-3 and AQ-4 exhibited the most evidence that they are able to stop the aggregation (or trap the aggregates) at the membrane interface. AQ-4 showed the highest ratio of oligomers by SEC, a constant random-coil signal from CD as well as the same 5 residues maintaining signal intensity in samples with and without membrane. This could indicate that even though AQ-4 interacts with the membrane, it is also able to directly interact with Aβ_40_. BF3 showed similar results as that observed for AQ-4. It showed no aggregation by ThT, maintained observable NMR signal intensity but induced an overall loss in intensity for every residue and also showed smaller species by SEC; this observation could indicate that while BF3 does not allow Aβ_40_ to aggregate, some Aβ_40_ population is still binding to the membrane surface. While DHQ-1 DHQ-2 and THQ-1 are less membrane active, they are still novel scaffolds for the inhibition of Aβ_40_. Although DHQ-1 and DHQ-2 showed some differences in activity, they may be a good starting point for developing derivatives. Given their similar architecture and sites for potential for synthesis, they could be used in an interesting study for structure activity relationship (SAR) analysis. The initial investigation and subsequent rule out of THQ-1 demonstrates the need for deep characterization for amyloid and small molecule interactions.

**Table 1.**
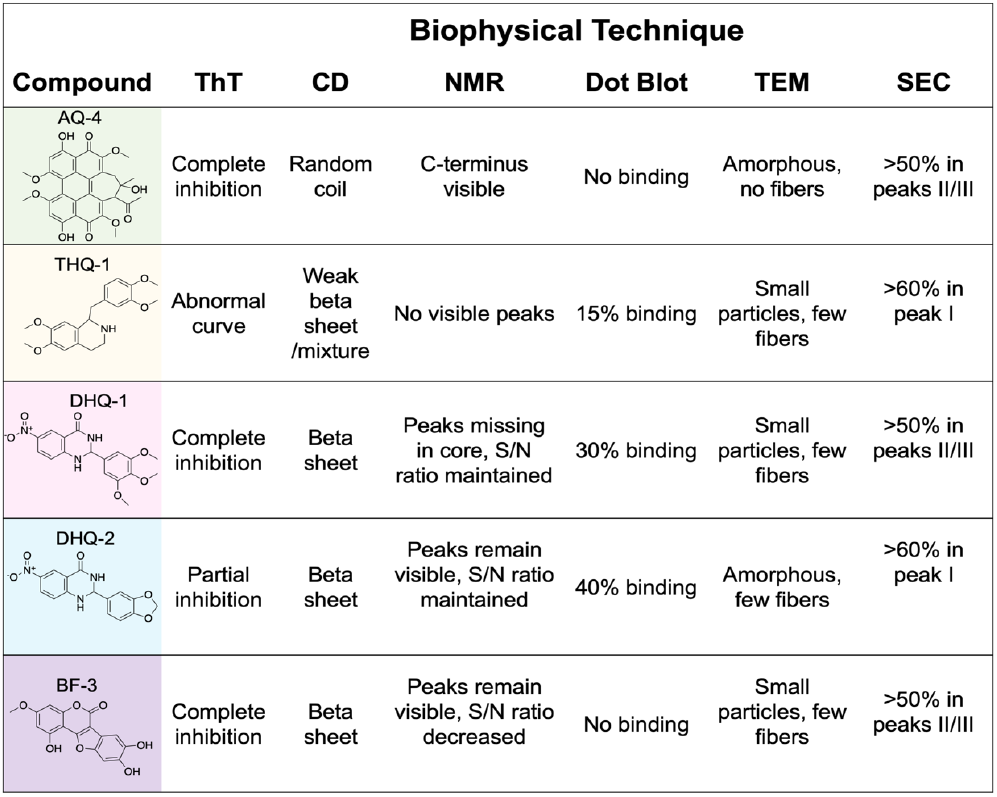
Summary of findings from the biophysical characterization of amyloid inhibition by the 5 compounds.

Given the similarity of membrane activities of AQ-4 and BF-3, it is possible that this may be due to their planar structure and lack of free rotation among the aromatic groups. However, many compounds initially investigated as part of the 21 primary hits produced prominent fibers; so clearly there must be unique properties possessed by these two compounds and their interplay between Aβ_40_ and the membrane which render their inhibitory activities. With the presented results as a starting point, NMR would be a robust tool to further investigate the structure of the Aβ_40_ compound structure as well as in conjunction with the membrane.

## Conflicts of interest

There are no conflicts to declare.

## Acknowledgments

This study was supported by the NIH (AG048934 to A.R.)) and the Michigan Alzheimer’s disease center MADC (M.I.I). We thank Steve Vander Roest and the University of Michigan Center for Chemical Genomics HTS core.

