## Supporting information for "High Throughput Screening at the Membrane Interface Reveals New Inhibitors of Amyloid-β"

### Materials and Methods

#### Screening Protocol

A library of 1,800 compounds were selected from Pilot Prestwick, Navigator Pathways and Maybridge libraries from the Center of Chemical Genomics (CCG) from the University of Michigan High Throughput Screening core facility. Thioflavin-T was selected as the primary read out for the screen and was used at 20 μM final concentration. All reagents were kept on ice until dispensing onto the plate. The screening was performed using $A\beta_{40}$ expressed in *E.coli* and purified as described elsewhere[1] and was used at a final concentration of 10 μM. Large unilamellar vesicles (LUVs) of a 7:3 mixture of 1,2-dioleoyl-sn-glycero-3-phosphocholine (DOPC) and 1,2-dioleoyl-sn-glycero-3-phospho-(1'-rac-glycerol) (DOPG) were prepared from chloroform stocks and made into lipid films by evaporating chloroform under nitrogen gas and then by lyophilization. Films were then rehydrated into buffer and then extruded to a size of 100 nm and were used at a final concentration of 500 μM. lipids and extruder were purchased from Avanti polar lipids. A buffer of 20 mM Phosphate 50 mM NaCl at pH 7.4 was used in all experiments. The assay was performed in 384 polystyrene plates from corning (product number 3640) with a non-binding surface. Each well was equipped with a single 0.5 mm sterile glass bead that will be added using a custom made bead dispenser from LabTIE, in order to act as a miniature stir rod to overcome the surface tension to ensure sufficient mixing during shaking.[2] Buffer was initially loaded into plates at a volume of 15 μL before adding the compounds dissolved in DMSO at a final concentration of 100 μM using a Biomek FX liquid handling automatic workstation. The remaining buffer, ThT, LUVs and $A\beta_{40}$ were added using Plates Thermo Fischer Mutli Drop Combi to a final volume of 30 μL and then sealed. Initial fluoresce readings were done from each plate at 440ex/480em using a Perkin Elmer 2014 Envision Multilabel plate reader on bottom read setting. After the initial reading, plates were moved to a mutilplate shaker kept inside an incubator set at 37 °C and were shaken for 24 hours at 750 rpm. After 24 hours of heating and shaking the plates fluorescence intensity was measured again. Each compound was added to duplicate plates in order to rule out false positives and to overcome the reproducibility issues among wells often seen with amyloid aggregation assays. After initial hits were identified, the top 40 hits were validated via the same protocol as described; however the compounds were titrated at 8 concentrations between 2-100 μM using TTP Labtech's Mosquito nanolitre hit picking system. From the concentration response curves (CRC) 21 compounds were selected based on the generated IC50 values and as well as removing compounds known to be PAINS. After the 21 compounds were selected, fresh compounds were bought from manufactures as shown in Table S1 and dissolved in DMSO. Full kinetic ThT profiles were obtained for each compound at 100, 50 and 20 μM concentrations. Reagents were prepared in the same manner as done in the screen but with using multichannel pipettes in place of robotic handling. Samples were plated in quadruplicates on 384 well plates, maintained at 37 °C and subject to continuous, slow orbital shaking. Fluorescence readings were taken on a Biotek Synergy 2 microplate reader and were read from the bottom with an excitation wavelength of 440 nm (30 nm bandwidth) and an emission wavelength of 485 nm (20 nm bandwidth) at three-minute intervals.

#### Dot Blot Assay

2 μL from the 50 μM compound containing ThT assay wells were spotted onto the nitrocellulose membrane in duplicate. Wells containing only compound and the LUVs were also spotted for control. Non-specific binding sites were blocked by soaking in 5% BSA in TBS-T for 30 min at RT. The primary Anti-Amyloid Fibrils OC Antibody at a 1:2000 dilution in 5% BSA in TBS-T

for 30 min at RT. The membrane was washed with TBS-T (3 x 5 min). Rabbit secondary antibody at a 1:1000 dilution was incubated for 30 min at RT. The membrane was washed with TBS-T (1 x 15 min and 2 x 5 min), and then incubated with ECL reagent for 1 min and sealed. Immediately after ECL reagent incubation, membranes were exposed using X-ray film in a dark room for 30 seconds. Images and intensities of spots were processed using ImageJ software.

**Transmission Electron Microscopy**

Samples used in TEM analysis were taken directly from ThT experiments on the wells of 5 equivalents of compound after 24 h incubation. Glow discharged grids (Formar/carbon 200 mesh, Electron Microscopy Sciences, Hatfield, PA, U.S.A.) were treated with samples (7 µL) for 2 min at room temperature. Excess buffer was removed via blotting and then washed three times with dd$H_2O$. Each grid was then incubated with uranyl acetate staining solution (1% w/v in dd$H_2O$, 7 µL) for 1 min, and excess stain was blotted away. Images from each sample were taken on a JEOL 1400-plus TEM (80 kV) at varying magnifications.

**Small molecule incorporation into lipids**

Each of the 5 compounds were dissolved into chloroform stocks and then added to lipid mixtures in chloroform at a 10:1 lipid to compound molar ratio. The lipid films were made as described above for a final concentration of 50 µM compound within 500 µM lipids. The compound containing LUVs were extruded and then checked for incorporation using UV-Vis measurements. Extruded solutions appeared clear and maintained clarity for up to 7 days on the bench top.

**Circular Dichroism**

Single point CD measurements were carried out at 25 µM $A\beta_{40}$ concentration in the presence of 500 µM lipids loaded with 50 µM of compound. CD measurements were performed on a JASCO J-1500 CD Spectrometer using a 0.1 cm path-length cell. Spectra were acquired at 25 °C using a bandwidth of 2 nm, a scan rate of 200 nm/min, and averaging spectra over 10 scans with a smoothing factor of 13. Measurements were taken immediately after mixing the sample (t = 0 h) and after various time points as mentined in the main text and figure captions.

**NMR experiments**

2D $^1H/^{15}N$ SOFAST-HMQC NMR spectra were recorded on a Bruker 800 MHz NMR spectrometer equipped with a 5mm triple resonance inverse detection TCI cryoprobe. NMR samples were prepared in 20 mM phosphate, 50 mM NaCl buffer (pH 7.4) containing 10% $D_2O$ at 298.2 K. Freshly dissolved $^{15}N$-labeled $A\beta_{40}$ was mixed with liposomes containing 500 µM of lipids and 50 µM of incorporated compound, as well as in the absence of lipids with the compounds dissolved in deuterated DMSO at 50 µM. All NMR samples were stored at 37 °C during the time-lapse measurements. Each 2D NMR spectrum was acquired using 256 $t_1$ increments, 16 scans, 16 dummy scans and a recycle delay of 0.2s. NMR data were processed using TopSpin 3.5 (Bruker) and analyzed using Sparky 3.114.

**Size exclusion chromatography**

NMR samples were used for SEC characterization. 200 μL of each sample was loaded using Superdex 75 Increase LS Tricorn 10/300 column operated on an AKTA purifier (GE Healthcare, Freiburg, Germany) read at 214 nm with a flow rate of 1 mL/min.

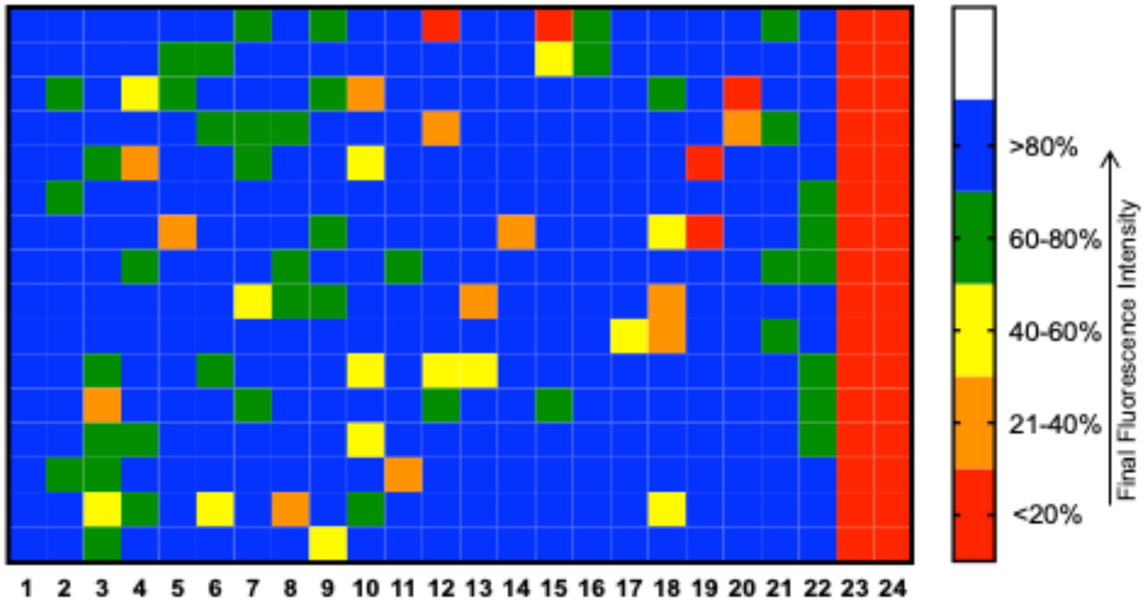


**Figure S1.** Representative data of final ThT fluorescence intensity of a 384 well plate from the screen. Columns 1 and 2 were negative controls and columns 23 and 24 were positive controls. Each individual well had a unique compound outside of the controls. Wells with lowest intensity (red or orange) were further investigated in this study.

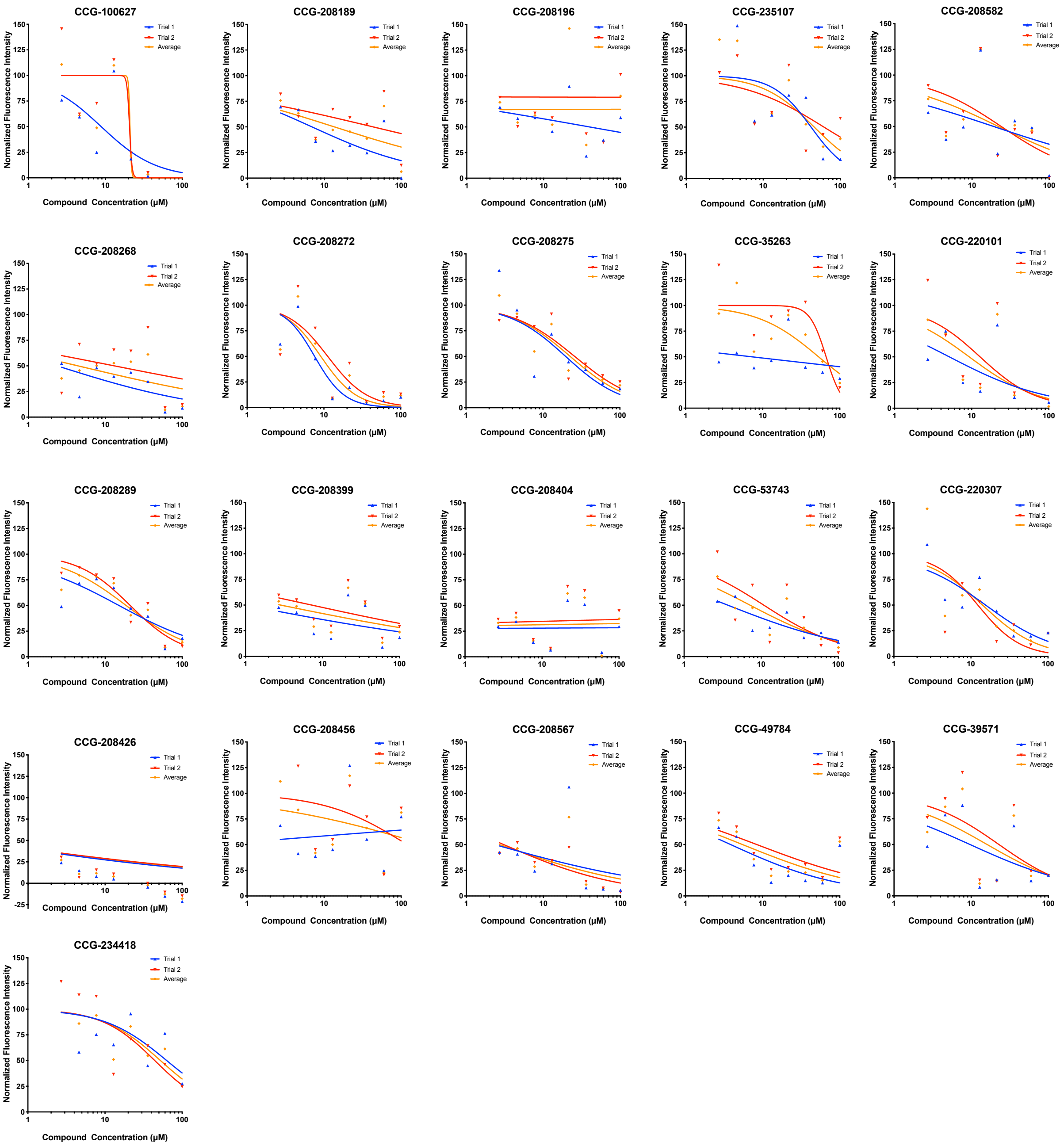


**Figure S2.** Concentration response curves for the 21 compounds selected further investigation. The concentrations of compounds ranged from 2-100 μM.

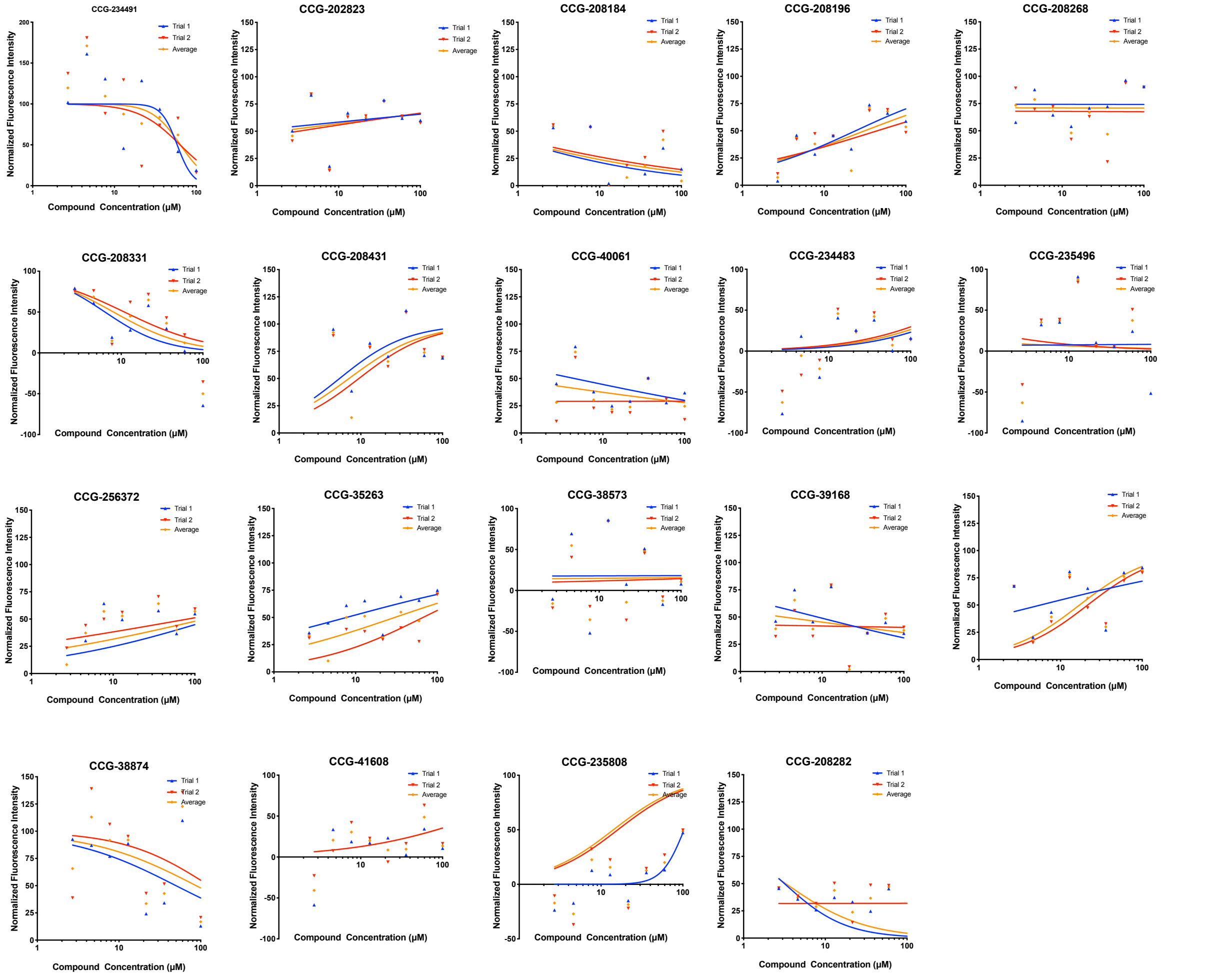


**Figure S3**. Concentration response curves for compounds not selected for further testing. Compound concentrations ranged from 2-100 μM.

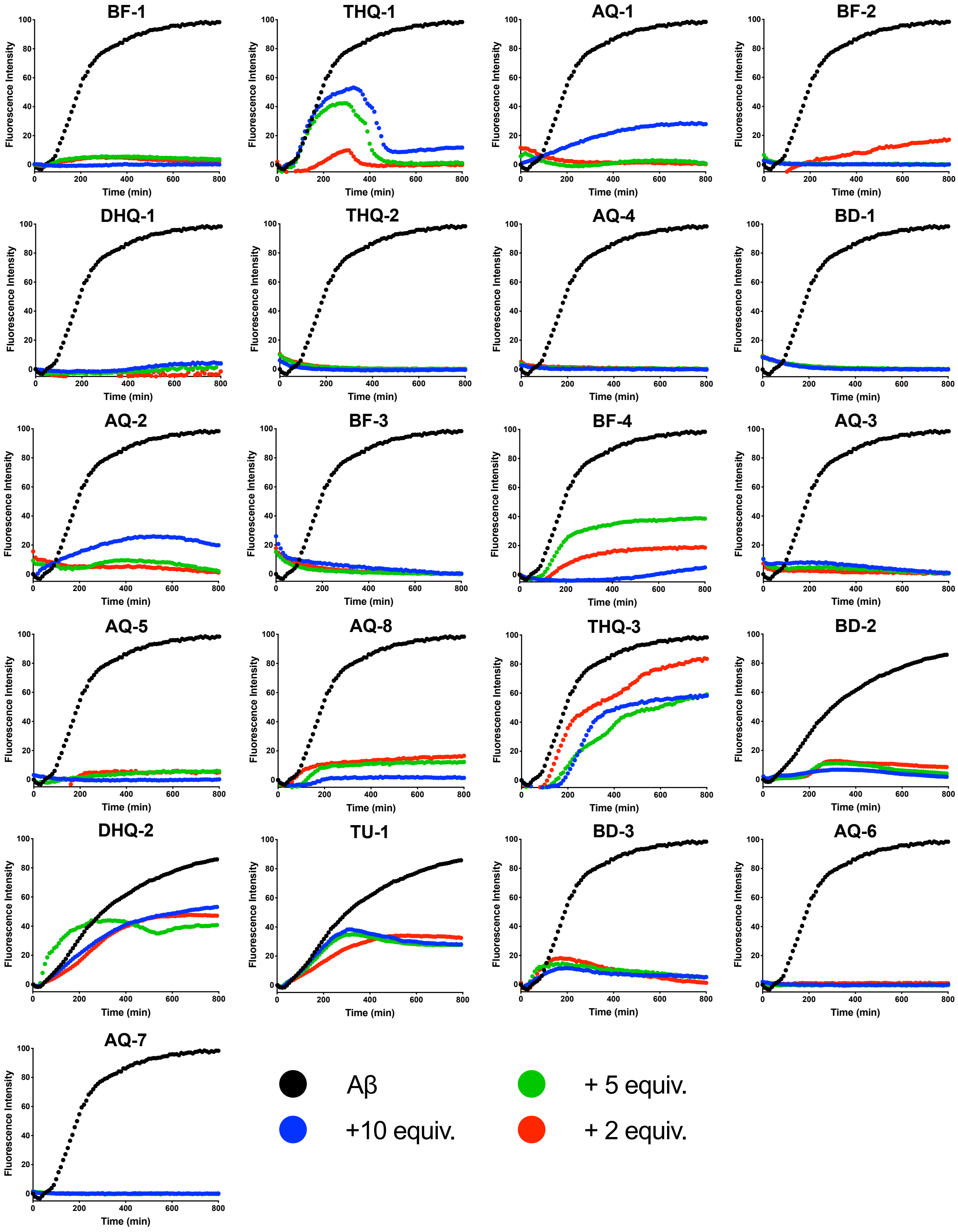


**Figure S4.** ThT fluorescence kinetics for the 21 selected compounds from CRC testing. 10 μM $A\beta_{40}$ was mixed with varying concentrations of small molecule (10, 5, & 2 equivalents) and 500 μM LUVs in 20 mM phosphate 50 mM NaCl buffer at 37 °C with slow shaking.

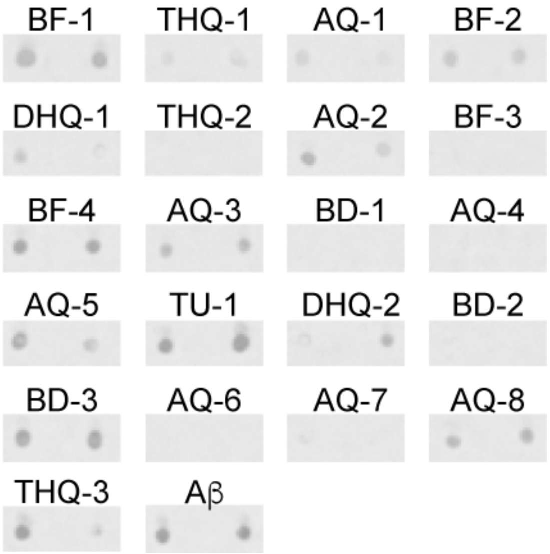


**Figure S5.** Dot blot images of samples after the ThT experiments using the OC antibody. Wells containing 10 μM of $A\beta_{40}$ + LUVs with 5 equivalents of the small molecule used for blotting.

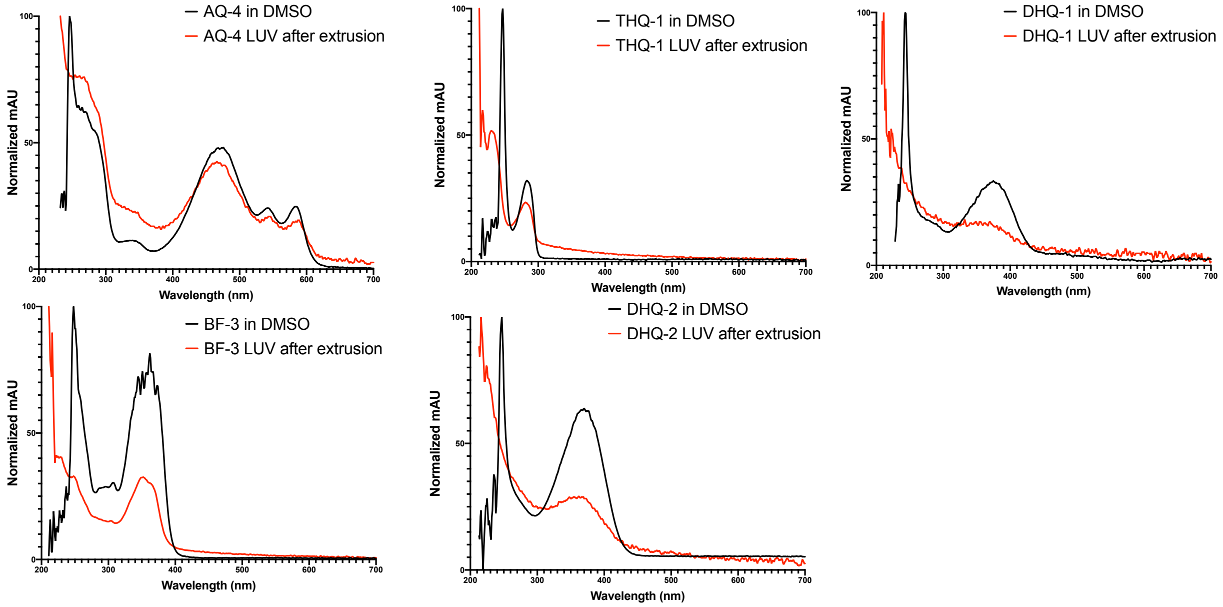


**Figure S6.** UV-Vis spectra of AQ-4, THQ-1, BF3-, DHQ-1, and DHQ-2 in DMSO (black) and incorporated into LUVs after extrusion (red). Molar ratio of lipids to compound was 10:1.

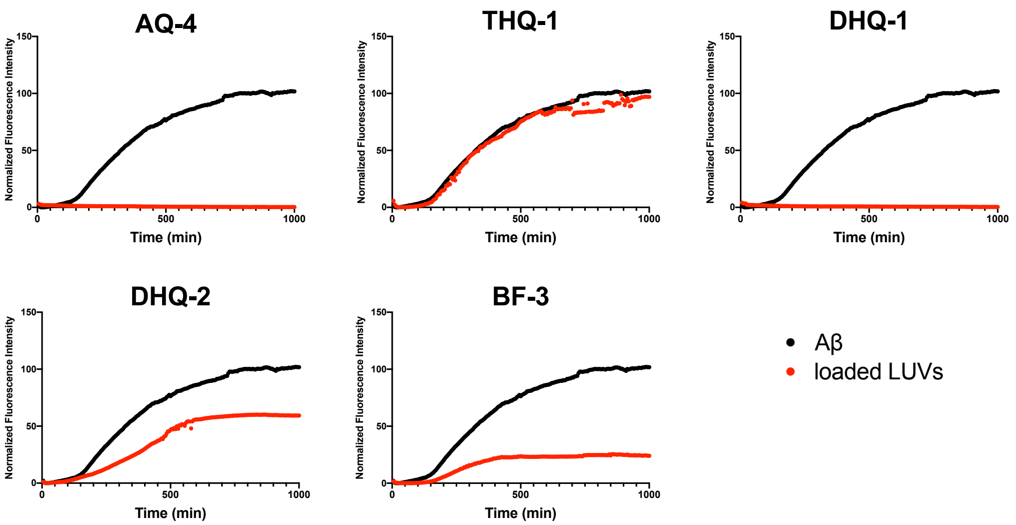


**Figure S7**. ThT fluorescence kinetics of 10 μM $A\beta_{40}$ (black) and $A\beta_{40}$ + 500 μM of compound loaded LUVs (red) at a 10:1 lipid:compound molar ratio.

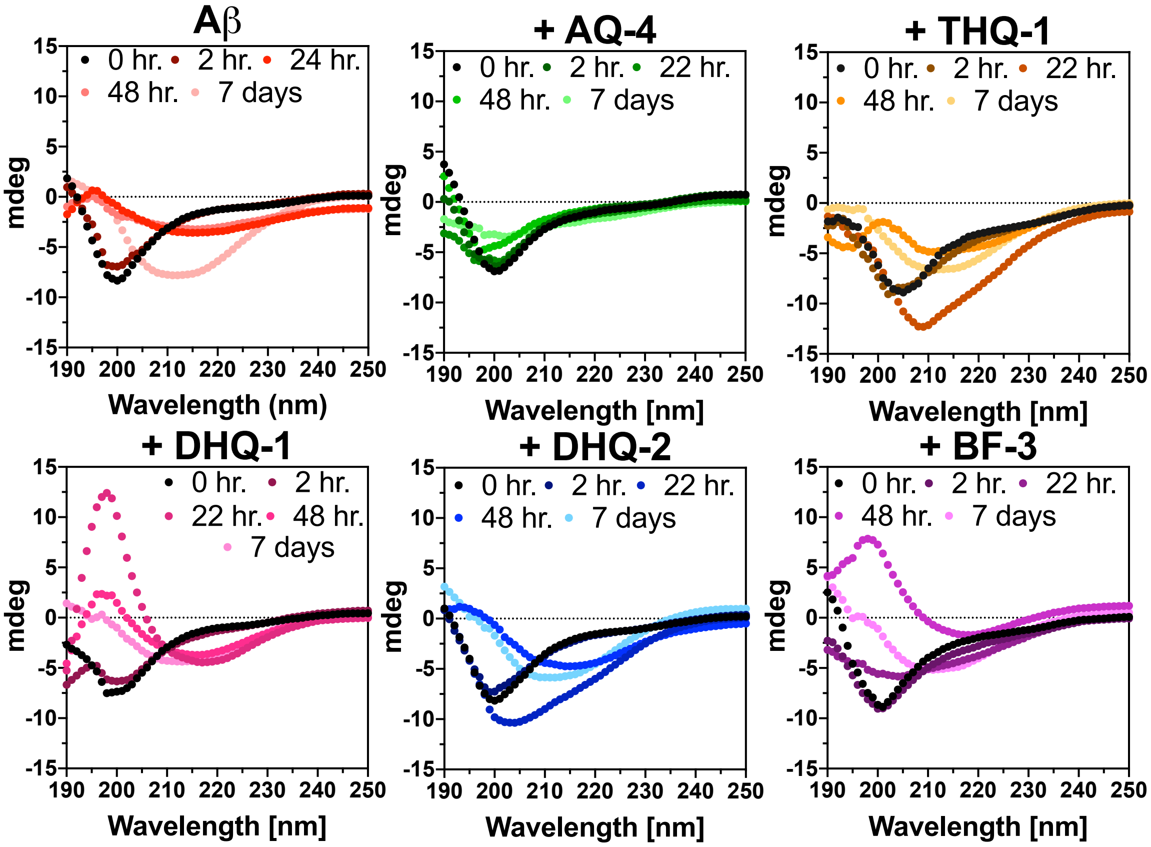


Figure S8. time course CD spectra of 25 μM Aβ40 in the presence of 500 μM of LUVs with 50 μM loaded compound LUVs. 48 hour trace shown in Figure 2C

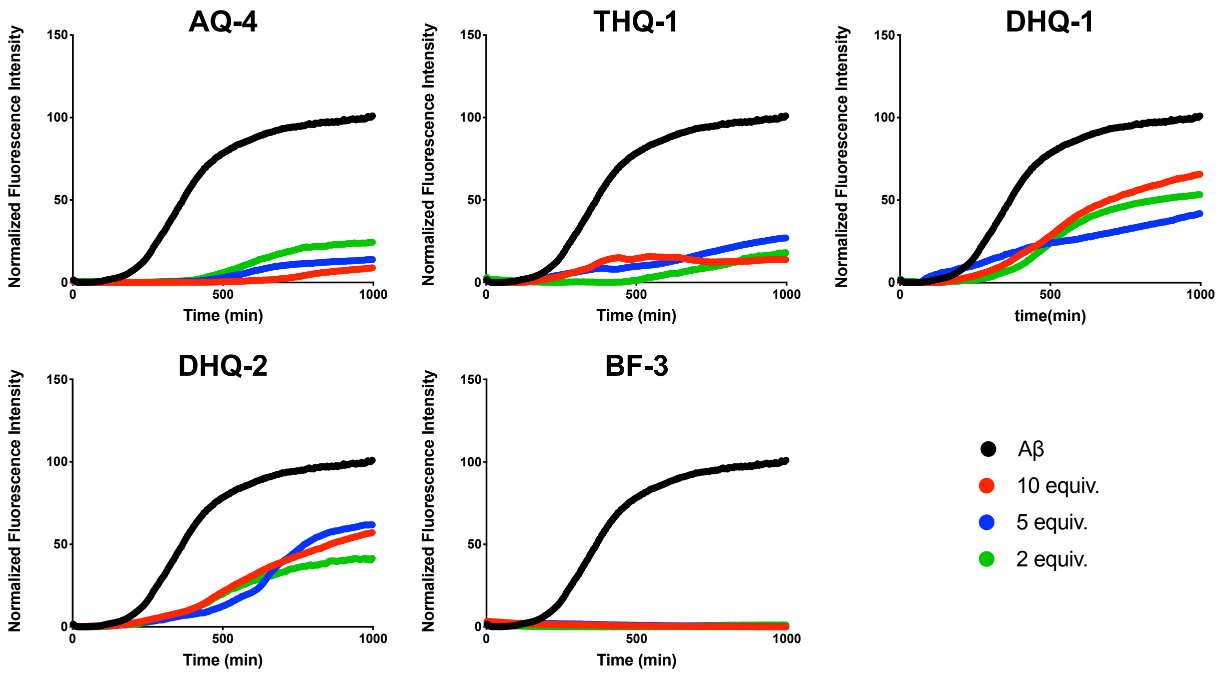


**Figure S9.** ThT fluorescence kinetics for 10 μM $A\beta_{40}$ (black) and varying equivalents of compound in the absence of lipids.

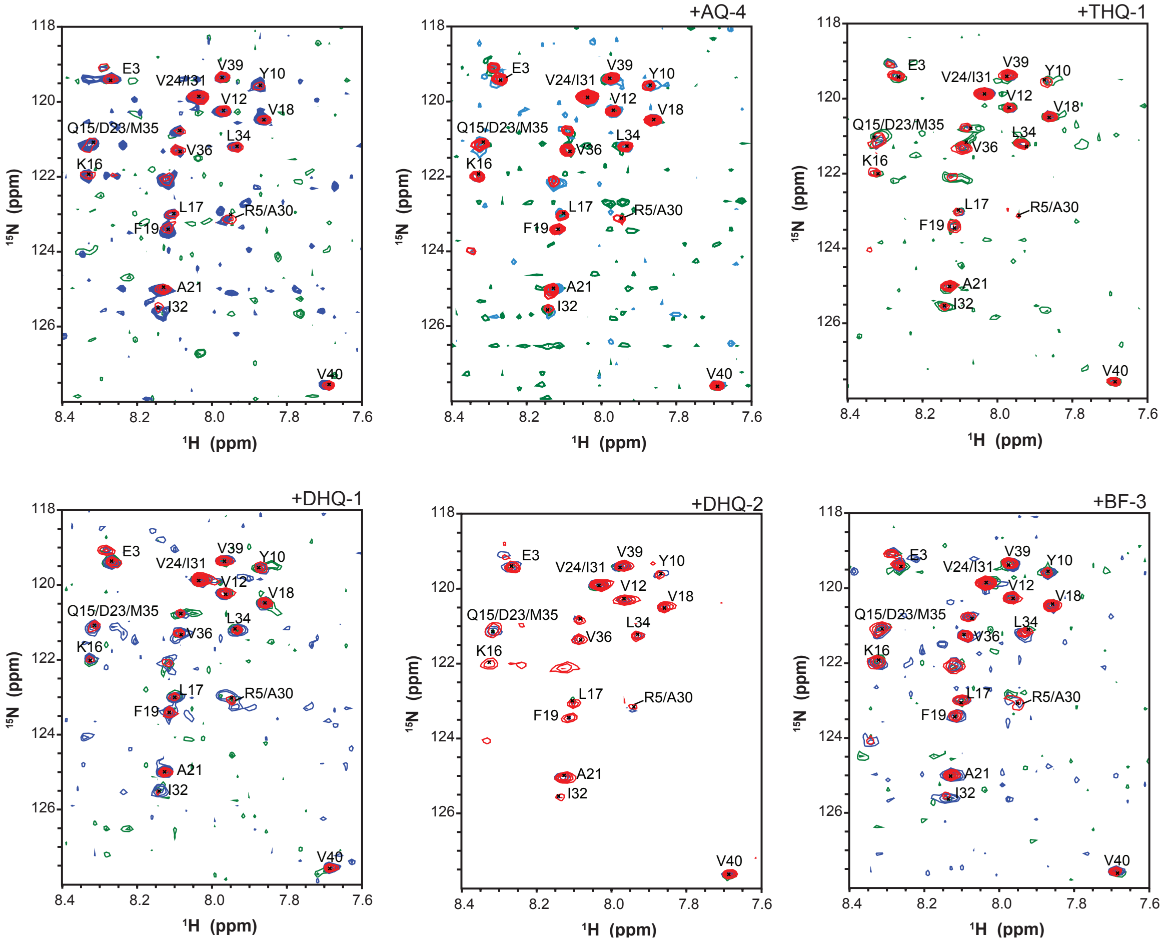


**Figure S10**. 2D SOFAST-HMQC NMR spectra of 25 μM $^{15}N$-labeled $A\beta_{40}$ in the absence of lipids with 2 equivalents of each compound at 0 hours (red), 24 hours (blue), and 96 hours (green).

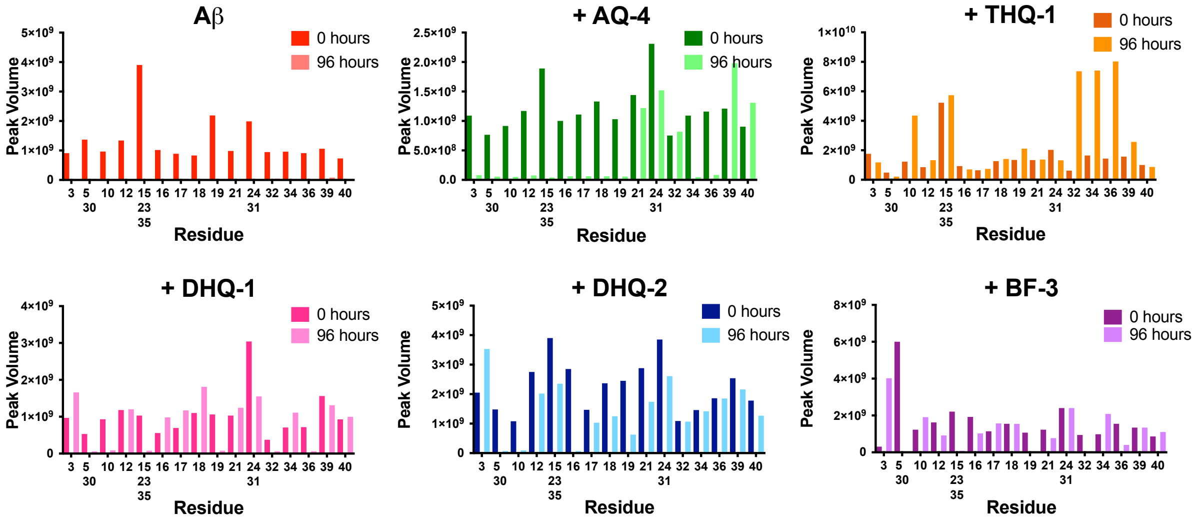


**Figure S11.** Peak volume measured from 2D SOFAST-HMQC NMR spectra of 25 μM [15]N-labeled $A\beta_{40}$ in the absence of lipids with 2 equivalents of each compound at 0 hours and 96 hours.

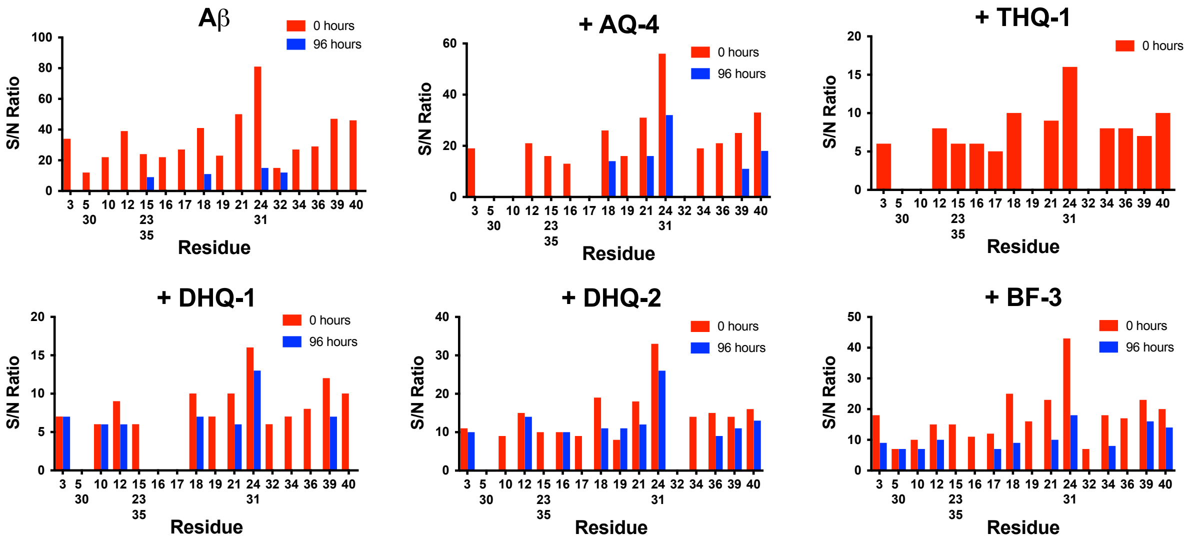


**Figure S12.** Signal-to-noise ratios measured from 2D SOFAST-HMQC NMR spectra of 25 μM $^{15}N$-A$\beta_{40}$ in the presence 500 μM of LUVs containing 50 μM loaded compound at 0 and 96 hours.

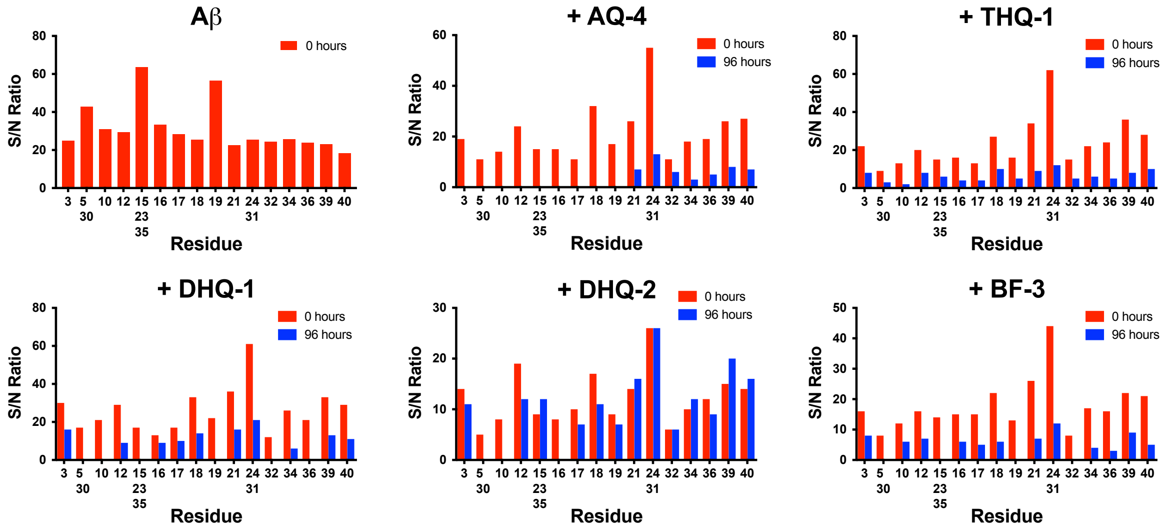


**Figure S13**. Signal-to-noise ratios measured from 2D SOFAST-HMQC NMR spectra of 25 μM $^{15}$N-A$\beta_{40}$ in the absence of lipids with 2 equivalents of each compound at 0 and 96 hours.

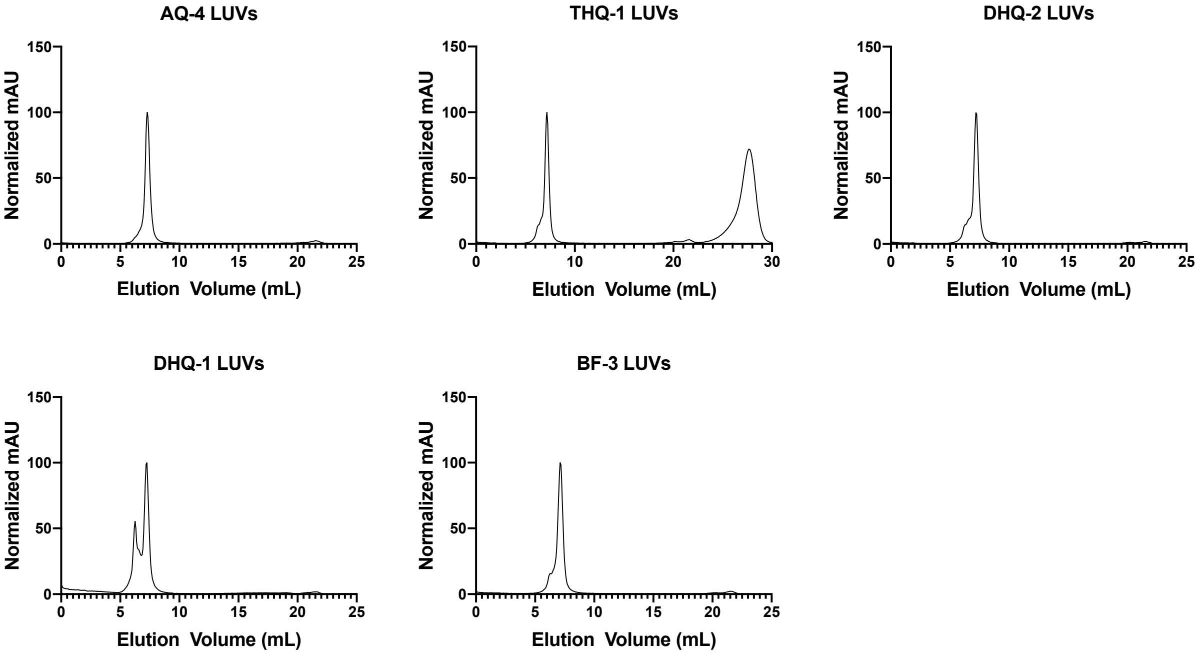


**Figure S14.** SEC profiles of 100 nm LUVs loaded with small molecules at a 10:1 lipid:compound molar ratio.

| CCG Code | Average IC50 | Name (IUPAC or Common) | Supplier |
|---|---|---|---|
| CCG-208404 | 15 | Maritimein | Carbosynth |
| CCG-49784 | 5.4 | 1-(3,4-dimethoxybenzyl)-6,7-dimethoxy-1,2,3,4-tetrahydroisoquinoline hydrochloride | Sigma Aldrich |
| CCG-208399 | 2.86 | Shikonin beta | Sigma Aldrich |
| CCG-208567 | 2.7 | Dihydrotanshinone | Molport |
| CCG-53743 | 7.2 | 2-(1,3-benzodioxol-5-yl)-6-nitro-1,2,3,4-tetrahydroquinazolin-4-one | Molport |
| CCG-220101 | 9.2 | Apomorphine hydrochloride hemihydrate | Sigma Aldrich |
| CCG-208272 | 9.114 | Shikonin | Sigma Aldrich |
| CCG-208289 | 18.7 | Wedelolactone | Molport |
| CCG-208275 | 22.6 | Tanshinone IIA | Sigma Aldrich |
| CCG-220307 | 14.15 | Oxytetracycline dihydrate | Sigma Aldrich |
| CCG-100627 | 20 | Piceatannol | Molport |
| CCG-208582 | 21 | Hypocrellin A | Carbosynth |
| CCG-208456 | 207 | Aloe-emodine | Molport |
| CCG-235107 | 50 | 1-(3,4-dimethoxyphenyl)-3-(3-methoxyphenyl)thiourea | Molport |
| CCG-234418 | 50 | 6-nitro-2-(3,4,5-trimethoxyphenyl)-2,3-dihydro-1H-quinazolin-4-one | Molport |
| CCG-39571 | 16 | Levodopa | Sigma Aldrich |
| CCG-208268 | 4 | Rosmarinic acid | Sigma Aldrich |
| CCG-208196 | 50 | Doxorubicin | Molport |
| CCG-208189 | 13 | Daunorubicin | Molport |
| CCG-35263 | 53 | Emodin | Sigma Aldrich |
| CCG-208426 | 0.18 | Laudanosoline | Molport |

Table S1. Information on the 21 primary hits selected form the screen.

| Common or IUPAC Name | Abbreviation | Structure |
|---|---|---|
| Maritimein | BF-1 | 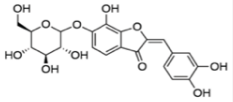 |
| 1-[(3,4-dimethoxyphenyl)methyl]-6,7-dimethoxy-1,2,3,4-tetrahydroisoquinoline hydrochloride | THQ-1 | 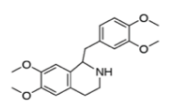 |
| Shikonin beta | AQ-1 | 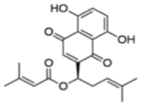 |
| Dihydrotanshinone | BF-2 | 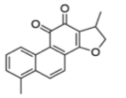 |
| 2-(1,3-benzodioxol-5-yl)-6-nitro-1,2,3,4-tetrahydroquinazolin-4-one | DHQ-1 | 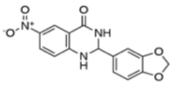 |
| Apomorphine hydrochloride | THQ-2 | 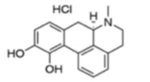 |
| Shikonin | AQ-2 | 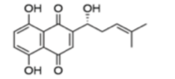 |
| Wedelolactone | BF-3 | 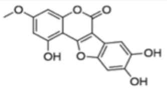 |
| Tanshinone IIA | BF-4 | 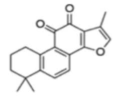 |
| Oxytetracycline | AQ-3 | 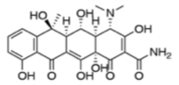 |
| Piceatannol | BD-1 | 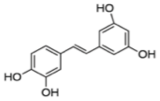 |
| Hypocrellin A | AQ-4 | 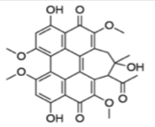 |
| Aloe-emodine | AQ-5 | 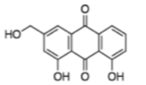 |
| 3-(3,4-dimethoxyphenyl)-1-(3-methoxyphenyl)thiourea | TU-1 | 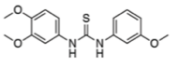 |
| 6-nitro-2-(3,4,5-trimethoxyphenyl)-2,3-dihydro-1H-quinazolin-4-one | DHQ-2 | 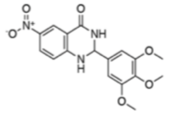 |
| Levodopa | BD-2 | 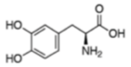 |
| Rosmarinic acid | BD-3 | 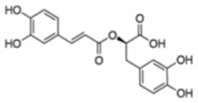 |
| Doxorubicin | AQ-6 | 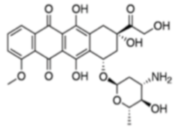 |
| Daunorubicin | AQ-7 | 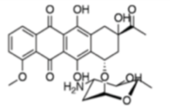 |
| Emodin | AQ-8 | 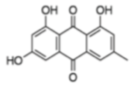 |
| Laudanosoline | THQ-3 | 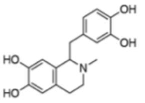 |

Table S2. Common name, given abbreviation, and structure of the 21 primary hits.
